# Acquisition of the L452R mutation in the ACE2-binding interface of Spike protein triggers recent massive expansion of SARS-Cov-2 variants

**DOI:** 10.1101/2021.02.22.432189

**Authors:** Veronika Tchesnokova, Hemantha Kulakesara, Lydia Larson, Victoria Bowers, Elena Rechkina, Dagmara Kisiela, Yulia Sledneva, Debarati Choudhury, Iryna Maslova, Kai Deng, Kirthi Kutumbaka, Hao Geng, Curtis Fowler, Dina Greene, James Ralston, Mansour Samadpour, Evgeni Sokurenko

**Affiliations:** University of Washington, Seattle, WA; ID Genomics, Inc., Seattle, WA; IEH Laboratories and Consulting Group, Seattle, WA; ARMADA (The Antibiotic Resistance Monitoring, Analysis and Diagnostics Alliance), Seattle, WA; Kaiser Permanente Washington (KPWA) and KPWA Research Institute, Seattle, WA

**Author notes:** Corresponding author: Evgeni V. Sokurenko.

## Abstract

The recent rise in mutational variants of SARS-CoV-2, especially with changes in the Spike protein, is of significant concern due to the potential ability for these mutations to increase viral infectivity, virulence and/or ability to escape protective antibodies. Here, we investigated genetic variations in a 414-583 amino acid region of the Spike protein, partially encompassing the ACE2 receptor-binding domain (RBD), across a subset of 570 nasopharyngeal samples isolated between April 2020 and February 2021, from Washington, California, Arizona, Colorado, Minnesota and Illinois. We found that samples isolated since November have an increased number of amino acid mutations in the region, with L452R being the dominant mutation. This mutation is associated with a recently discovered CAL.20C viral variant from clade 20C, lineage B.1.429, that since November-December 2020 is associated with multiple outbreaks and is undergoing massive expansion across California. In some samples, however, we found a distinct L452R-carrying variant of the virus that, upon detailed analysis of the GISAID database genomes, is also circulating primarily in California, but emerged even more recently.

The newly identified variant derives from the clade 20A (lineage B.1.232) and is named CAL.20A. We also found that the SARS-CoV-2 strain that caused the only recorded case of infection in an ape - gorillas in the San Diego Zoo, reported in January 2021 - is CAL.20A. In contrast to CAL.20C that carries two additional to L452R mutations in the Spike protein, L452R is the only mutation found in CAL.20A. According to the phylogenetic analysis, however, emergence of CAL.20C was also specifically triggered by acquisition of the L452R mutation. Further analysis of GISAID-deposited genomes revealed that several independent L452R-carrying lineages have recently emerged across the globe, with over 90% of the isolates reported between December 2020 – February 2021. Taken together, these results indicate that the L452R mutation alone is of significant adaptive value to SARS-CoV-2 and, apparently, the positive selection for this mutation became particularly strong only recently, possibly reflecting viral adaptation to the containment measures or increasing population immunity. While the functional impact of L452R has not yet been extensively evaluated, leucine-452 is positioned in the receptor-binding motif of RBD, in the interface of direct contact with the ACE2 receptor. Its replacement with arginine is predicted to result in both a much stronger binding to the receptor and escape from neutralizing antibodies. If true, this in turn might lead to significantly increased infectivity of the L452R variants, warranting their close surveillance and in-depth functional studies.

## Introduction

The recent emergence of mutational variants of SARS-CoV-2 (nCoV) around the globe suggests adaptive evolution of the virus, potentially affecting its transmissibility, infectivity, virulence and/or immune escape [1–4]. The primary target of current vaccines and monoclonal antibodies is the Spike protein which mediates viral attachment to and entry into host cells [5, 6]. Thus, emergence of variants with mutations in the Spike protein are of particular interest due to their potential for reduced susceptibility to neutralizing antibodies elicited by vaccination or prior infection.

The Spike protein is 1,273 amino acids long and is comprised of the N-terminal region S1 (amino acids 14-682) responsible for viral attachment to target cells via the ACE2 receptor, and the C-terminal region S2 (aa 686–1273) responsible for membrane fusion and cell entry [7]. Before fusion, S1 is cleaved from S2 in the cleavage region (aa 682-685). Antibodies against the ACE2 binding domain of S1 (Receptor-Binding Domain, RBD; aa 319-541) are considered to be critical in neutralizing nCoV [8–10]. Because of the important functional and antigenic properties of RBD, structural changes in this domain deserve special attention and have already been highlighted by such notorious RBD mutations as E484K (e.g. found in the ‘Brazil’ variant B.1.1.28) or N501Y (found in the ‘British’ B.1.1.7 variant and ‘South African’ B.1.351 variant) [1].

We evaluated the feasibility of determining mutational changes in the S1 region by PCR amplification of 541 bases fragment within and immediately downstream from the RBD coding region, followed by Sanger sequencing (see Methods). The amplicon included the gene region coding aa 414 to 583 of the Spike protein (for further: ‘region 414-583’), which includes the so-called receptor-binding ridge epitope (aa 417, 455, 456 and 470–490), the 443–450 loop epitope (aa 443–452 and 494–501) and the 570-572 loop of the so-called C-terminal domain 1 (CTD1) of S1, which is important in the interaction between the S1 and S2 regions. As test samples, we used 570 clinical oropharyngeal specimens collected during April-May from nCoV-positive patients at Kaiser Permanente Washington (51 samples) and nCoV-positive clinical samples submitted for testing to IEH Laboratories, Inc. (Bothell, WA), from four states (California, Washington, Arizona, Colorado, Minnesota and Illinois) that were split into a September-October collection group (85 samples) and a November-February collection group (434 samples). In the process of region 414-583 analysis, we noted a high prevalence of samples that carried nCoV variants with L452R mutation. This prompted us to perform an in-depth follow-up analysis of the L452R-carring nCoV strains in our samples and, then, their prevalence and clonal origin on the global scale by using publicly available nCoV genomes and analytical tools of GISAID and Nextstrain databases.

## Results

It was possible to amplify the 414-583 region and obtain high quality sequence from 99.8% of the specimens. A total of 58 of the samples had 1 to 2 changes from the corresponding region of the reference Wuhan-Hu-1 genome (NC_045512). All sequence changes in the region were single nucleotide polymorphisms (SNPs). Among the April-May samples, only one sample (2.0%) was found with a mutation. This mutation was of silent (synonymous) nature, i. e. without the change in amino acid content (**Table 1**). Among the September-October samples, four samples (4.7%) had single SNPs, including three silent mutations and one amino acid (nonsynonymous) mutation. Among the November-February samples, 53 (12.2%) had a total of 15 silent SNPs and 11 amino acid (nonsynonymous) mutations. Among the later samples, all multiple samples tested at the IEH Laboratories that were known to have been isolated from the same patient or submitted from the same collection site on the same day had the same mutational profiles and were considered as epidemiologically linked and considered as duplicated. Upon removal of the duplicates, 39 non-linked samples contained mutations in the 414-583 region, with 27 unique SNPs or SNP combinations.

**Table 1.** Mutations distribution across the test samples. No. samples column shows total number of samples from a site received same day. In bold – amino acid changes

| Time period | No. samples | Site | State | Region 1 (414-583) | Region 2 (1-250) |
| --- | --- | --- | --- | --- | --- |
| **April - May (N=51)** | 1 | KP | WA | c1425t |  |
| **September - October (N=85)** | 1 | 22 | CA | t1695c |  |
|  | 1 | 4 | CA | c1497t |  |
|  | 1 | 10 | CA | c1497t |  |
|  | 1 | 23 | PA | g1450a (**E484K**) |  |
| **November - February (N = 434)** | 1 | 3 | CA | a1290g |  |
|  | 1 | 3 | CA | t1326c |  |
|  | 1 | 20 | MN | t1338g |  |
|  | 1 | 6 | AZ | t1356g |  |
|  | 1 | 4 | CA | c1425t |  |
|  | 1 | un | un | t1458c |  |
|  | 1 | 15 | MN | t1506g |  |
|  | 2 | 18 | NM | c1524t |  |
|  | 1 | 13 | WA | c1626t |  |
|  | 1 | 7 | CO | c1770t |  |
|  | 1 | 2 | CA | t1355g (**L452R**) | g38t (**S13I**) + g456t (**W152C**) |
|  | 1 | 12 | WA | t1355g (**L452R**) | g38t (**S13I**) + g456t (**W152C**) |
|  | 2 | 4 | CA | t1355g (**L452R**) | g38t (**S13I**) + g456t (**W152C**) |
|  | 2 | 14 | WA | t1355g (**L452R**) | g38t (**S13I**) + g456t (**W152C**) |
|  | 1 | 17 | CA | t1355g (**L452R**) | g38t (**S13I**) + g456t (**W152C**) |
|  | 1 | 16 | CA | t1355g (**L452R**) | g38t (**S13I**) + g456t (**W152C**) |
|  | 2 | 5 | CA | t1355g (**L452R**) | g38t (**S13I**) + g456t (**W152C**) + del 141-144 LGVY |
|  | 1 | 4 | CA | t1355g (**L452R**) | g38t (**S13I**) + g456t (**W152C**) + c76t |
|  | 1 | 4 | CA | t1355g (**L452R**) + c1416t | g38t (**S13I**) + g456t (**W152C**) |
|  | 2 | 8 | WA | t1355g (**L452R**) + t1593c | g38t (**S13I**) + g456t (**W152C**) |
|  | 1 | 3 | CA | t1355g (**L452R**) + c1626t | g38t (**S13I**) + g456t (**W152C**) |
|  | 2 | 5 | CA | t1355g (**L452R**) |  |
|  | 2 | 4 | CA | t1355g (**L452R**) + t1715g (**T572I**) |  |
|  | 1 | 21 | GA | g1365t (**L455F**) |  |
|  | 1 | 4 | CA | c1387t (**P463S**) + t1503c |  |
|  | 1 | 7 | CO | c1409a (**T470N**) |  |
|  | 5 | 4 | CA | c1433a (**T478K**) |  |
|  | 1 | 19 | CA | c1433a (**T478K**) |  |
|  | 1 | 10 | CA | c1433a (**T478K**) + c1686t |  |
|  | 1 | 1 | MN | g1445t (**G482V**) |  |
|  | 1 | 1 | MN | g1450a (**E484K**) |  |
|  | 2 | 20 | MN | g1450a (**E484K**) |  |
|  | 1 | 13 | WA | g1450a (**E484K**) |  |
|  | 1 | 9 | WA | t1480c (**S494P**) |  |
|  | 1 | 7 | CO | g1696a (**G566S**) |  |
|  | 2 | 9 | WA | c1709t (**A570V**) |  |

Silent mutations were distributed across the 414-583 region, without clear clustering (**Figure 1**). In contrast, the amino acid changes were distributed non-randomly and, except for one mutation (P463S), were clustered within the main epitope regions of RBD (L452R, T470N, T478K, G482V, E484K and S494P) or in the 570-572 loop of CTD1 (A570V and T572I). All of the amino acid mutations had been reported previously and could be identified in nCoV sequences deposited to GISAID. Though the notorious N501Y mutation was not identified in our samples, five samples contained mutation E484K that is found in lineages from various countries and present in the ‘Brazil’ variant B.1.1.28. E484K is located in the receptor-binding ridge epitope and was shown to provide marked resistance to neutralizing antibodies in multiple studies [8, 11, 12].

**FIGURE 1:**
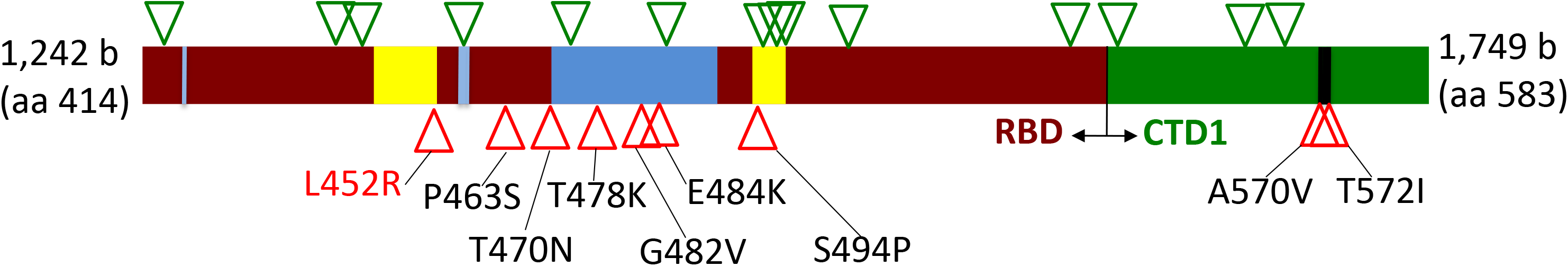
Distribution of silent (green triangles) and amino acid (red triangles) mutations across region 414-583 of the Spike protein. Dark red – receptor-binding domain (RBD); Green – C-terminal domain 1 (CTD1) of S1 Spike region; Blue – receptor-binding ridge epitope residues; Yellow – 443-450 loop epitope residues; Black – 570-572 loop residues.

The most frequent mutation found in the region 414-583 was by far L452R, occurring in 19 out of 39 (48.7%) total and 13 out of 26 (50.0%) non-linked samples, isolated in 9 out of 18 (50%) separate collection sites. Three samples with L452R also had separate silent changes and two samples (from the same site) had an additional amino acid change T572I. L452R was found mostly in samples (14) and sites (6) from California, though 5 samples from 3 sites came from Washington. L452R is part of the 443-450 loop epitope and is located on the edge of the receptor-binding motif of RBD formed by residues in direct contact with ACE2 [8, 13, 14]. Its occurrence appeared to be only sporadic in most of 2020 and has received significant attention only very recently due to the report of the California Department of Public Health released January 17, 2021 and a follow-up publication [15]. It was noted that there has been a recent sharp rise in isolation of nCoV variants with L452R across multiple outbreaks in California, accounting for more than a third of all isolates. According to the GISAID database, only six nCoV genomes with L452R were deposited in September-October 2020 (all from California), but since then additional genomes with the mutation have been deposited including 142 in November (95.7% from California), 488 in December (79.1%) and 619 in January 2021 (69.2%). This expansion has been linked to a single viral variant from clade 20C according to the Nextstrain nomenclature of nCoV (lineage B.1.429 according to the PANGO nomenclature of nCoV) and was designated as CAL.20C (20C/S:452R; B.1.429).

According to genomic analysis, CAL.20C has also been defined by 4 additional amino acid mutations, including two Spike protein mutations, S13I and W152C, located in the signal peptide and N-terminal domain, respectively [15]. Using an approach similar to our sequencing of the region 414-583, we amplified and sequenced the aa1-250 coding region in all samples with the L452R mutation. Both S13I and W152C mutations were found in 15 total (11 non-linked) samples suggesting their identity with CAL.20C. (Among those samples, one contained an additional silent mutation, while in two samples from one site a four-amino acid deletion (141-144 LGVY) was found.) Surprisingly, in 4 samples with L452R the additional mutations were absent. Those samples originated from two separate sites (two samples in each) in California. One sample pair carried the T572I mutation in the 414-583 region mentioned above. To determine how closely these S13I/W152C non-carrier L452R variants were related to the CAL.20C variant, full genome sequencing was possible performed on three of those samples and on four CAL.20C-like, S13I/W152C carrier samples. Based on the genome-wide analysis, all CAL.20C-like variants were in the same clade as the chosen reference CAL.20C strain (GISAID # 730092) isolated in September, 2020 (**Figure 2A**). In sharp contrast, the other L452R variants formed a distinct phylogenetic clade that is distant from CAL.20C, sharing none of the CAL.20C-specific mutations. In fact, further analysis established that those strains derived from a separate 20A clade, lineage B.1.232, indicating that the L452R mutation in this group of strains was acquired independently from CAL.20C. To distinguish from the recently described CAL.20C variant, we designated this novel L452R-carrying variant as CAL.20A (20A/S:452R/B.1.232).

**Figure 2.**
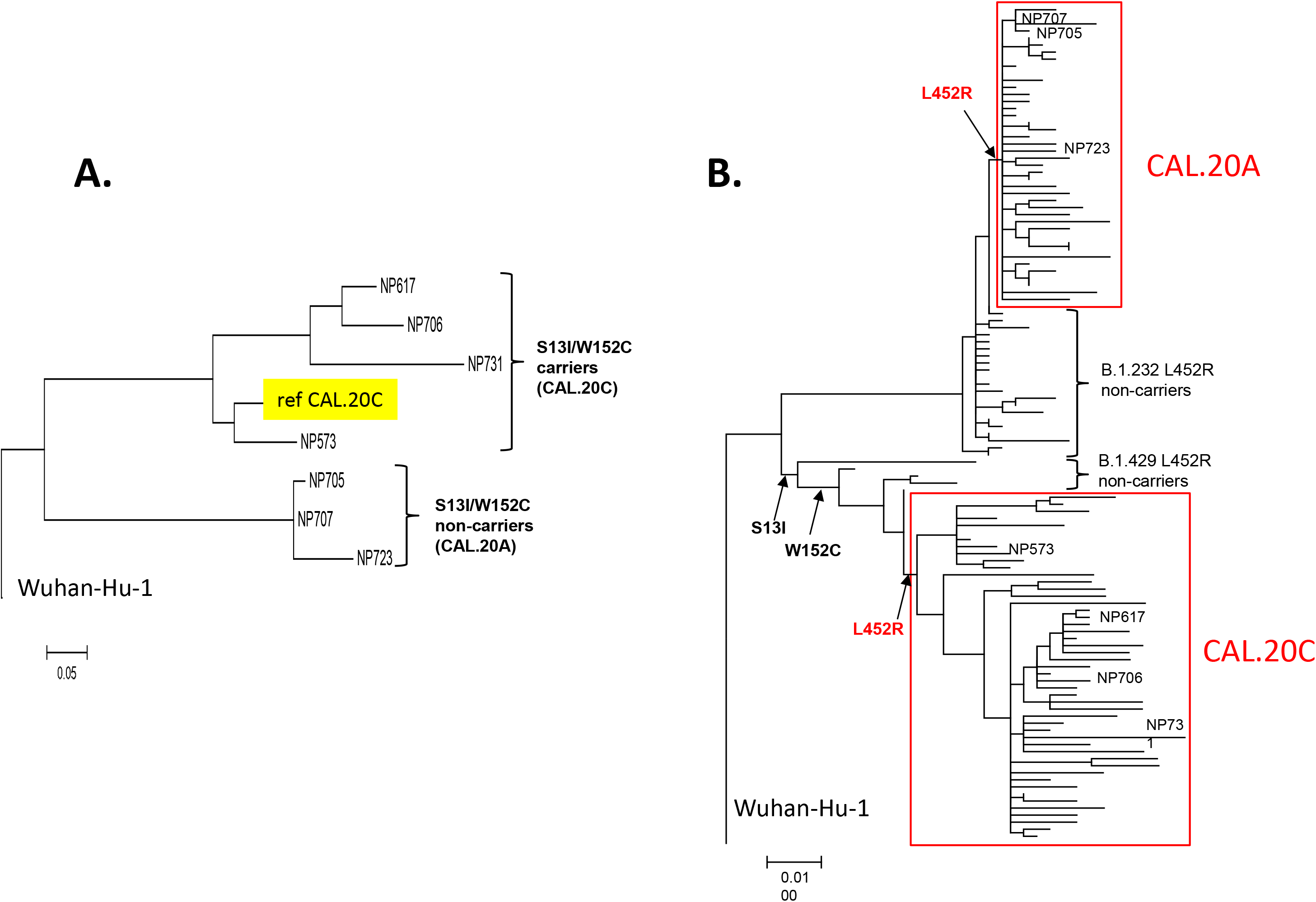
Phylogenetic trees of CAL.20A and CAL.20C genomes. A. nCoV strains identified in the tested samples. B. nCoV strains deposited into GISAID database.

Analysis of the GISAID database on February 19, 2021, revealed that the CAL.20A-including lineage B.1.232 contained a total of 559 deposited genomes, but only 54 of them (9.7%) contained the L452R mutation, all closely related to the CAL.20A strains identified here. The first deposition of an CAL.20A genome was made on November 23, 2020, from Baja California, Mexico (GISAID # 878300), with most of the CAL.20A genomes deposited in January-February 2021 (43 genomes; 79.6%). The vast majority of CAL.20A strains (74%) were isolated in the state of California, but also in 5 other states (WI, NM, UT, PA, AZ) as well as Canada, Mexico and Costa-Rica. Interestingly, one of the CAL.20A samples was derived from a gorilla in the San Diego Zoo (deposited to GISAID January 10; GISAID # 862722) – a highly publicized case of nCoV infection in apes [16].

To compare the clonal diversity of CAL.20A and CAL.20C strains, we build a phylogenetic tree of the January isolates of CAL.20A and 50 randomly selected genomes of CAL.20C also deposited in January 2021. We also included into the analysis some strains that we have identified as the most closely related to the L452R variants, but without this mutation. Based on the shorter branches overall, the CAL.20A cluster appeared to be less diverse than the CAL.20C cluster (**Figure 2B**). The pairwise difference in the number of silent, presumably neutrally accumulating mutations per genome was 2.56±1.41 and 8.22±4.19 mutations, respectively (P<.01), indicating that CAL.20A has emerged much more recently than CAL.20C.

In contrast to CAL.20C, L452R was the only omnipresent amino acid mutation in the Spike protein of CAL.20A, relatively to the Wuhan-Hu-1 reference strain. Other mutations, like T572I in our samples, were found only sporadically and in few strains (with most mutations in the S2 region). In fact, L452R was the only amino acid mutation in the entire genome that separated CAL.20A from the closest non-L452R strains within the B.1.232 lineage (GISAID # 636127, **Figure 2B**). Thus, acquisition of L452R appears to be the primary evolutionary event that led to emergence of CAL.20A. Though it was originally reported that, besides L452R, all CAL.20C strains carry 4 more specific amino acid mutations, including S13I and W152C in the Spike protein, we found that few closely related strains in the B.1.429 lineage carry either S13I alone (GISAID # 977963) or both S13I and W152 but not L542R (GISAID # 847642 and # 977918, **Figure 2B**). Thus, the Spike mutations in the CAL.20C-contaning lineage were acquired sequentially with the L452R acquired most recently, triggering the current massive clonal expansion of CAL.20C.

Examination of the Nextstrain and GISAID databases on February 18, 2021, revealed that, besides CAL.20A and CAL.20C, there are a total of 410 nCoV genomes that contain the L452R mutation which has been acquired by at least 5 separate lineages – A.21, A.2.4, B.1.1.10, B.1.1.130 and C.16 within different clades, ranging from 2 to 213 strains each (**Figure 3**). The strains were isolated from over 20 countries across all continents, with no apparent dominance of any geographic area. In one deposited strain, from lineage B.1.74, an L452Q mutation was present instead of L452R. There is a clear temporal trend in the number of deposited genomes, with 2 genomes only from July-September, 28 from October-November, 127 from December 2020 and 238 from January 2021. Thus, the L452R mutation has been acquired independently in a variety of clonally diverse nCoV strains, with 92.6% of L452R strains reported after November 2020, indicating a very recent emergence of all those lineages.

**FIGURE 3.**
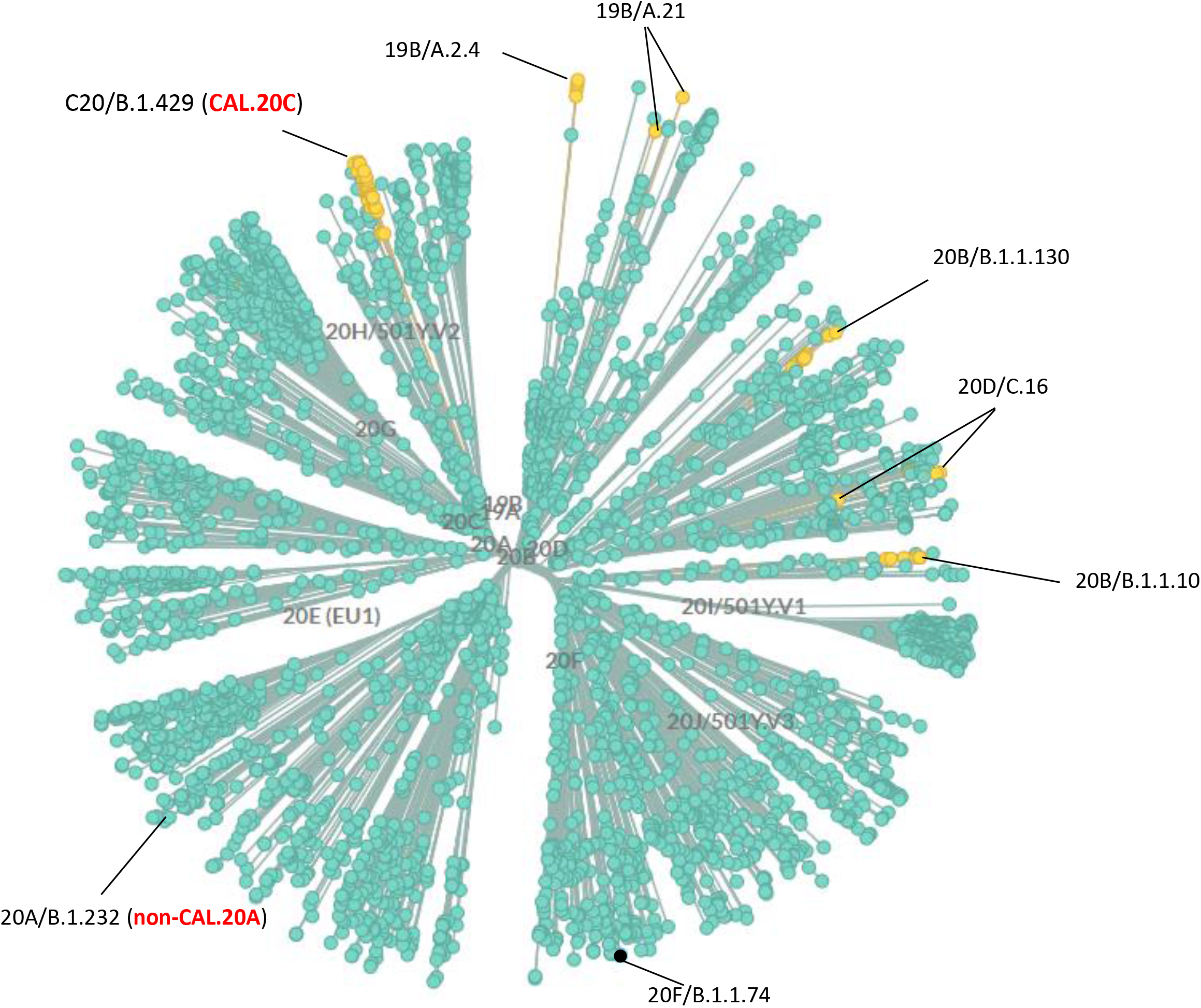
Nextstrain Unrooted cladogram phylogenetic tree of SARS-CoV-2 genetic variants (as of February 15 2021). Nodes in turquoise – L452; in yellow – R452; in black – Q452. Nomentlature: Nextstrain clade/PANGO lineage. At the time of analysis, CAL.20A strains were not found in the Nextstrain database and only one B.1.232 (non-CAL.20A) was found.

## Discussion

Taken together, our results show that two independent nCoV variants recently emerged in the state of California that carry the L452R mutation in the Spike protein, the already defined and currently dominant CAL.20C [15] as well as the more recently emerged CAL.20A identified here. The fact that, according to our analysis, emergence of both CAL.20A and CAL.20C was triggered by the L452R mutation alone provides direct evidence for the adaptive significance of this mutation specifically and, also, creates a potential opportunity to isolate and functionally compare naturally occurring isogenic variants of nCoV with and without L452R.

On the other hand, it is possible that the lack of other mutations specific to CAL.20A or shared with CAL.20C could be the reason why the former variant has not undergone as extensive expansion as CAL.20C. This would indicate (and availability of CAL.20A should help to affirm) that other mutations in CAL.20C might enhance the adaptive impact of L452R, i.e. that the genomic background of L542R plays a significant role as the target of positive selection. However, it is also possible that CAL.20A is at least as fit as CAL.20C, but has not expanded as broadly because it emerged much more recently and after CAL.20C has underwent extensive expansion in the same geographic niche area. Co-circulation of CAL.20A and CAL.20C in the same area provides a unique opportunity to study the interplay between variants in space and time in demographically diverse but interconnected communities and patient populations.

According to the public databases, in addition to the two California strains the L452R mutation has been acquired at this point by at least half a dozen independent lineages across multiple countries and continents. Though detailed examination of the timing, geography and genomic background of L452R emergence in different lineages is beyond the scope of this study, such repeatedly emerging hot-spot mutations typically indicate strong positive selection. Interestingly, it appears that the selection for L452R became especially strong very recently. This is possibly reflecting adaptive evolution of the virus in response to either the epidemiological containment measures extensively introduced in the fall of 2020 or a growing proportion of the population with immunity to the original viral variants, i.e. the reconvalescents and vaccinated individuals.

Though potentially an accidental event, isolation of CAL.20A from a gorilla at the San Diego Zoo is worthy of note. According to the sequence deposited in GISAID, the gorilla CAL.20A variant carries two additional SNPs, both in the ORF1ab non-structural protein 2 (nsp2), a silent c934t and non-synonymous c810t (T183I), but with several sequence stretches unfortunately missing. It is impossible to say at this point to what extent the isolation of the CAL.20A variant is connected to possibly distinctive biological properties of the strain and specifically the L452R mutation. However, it would be valuable to determine whether the occurrence of CAL.20A infection in the gorilla is due to specific features of CAL.20A with regard to viral transmissibility, infectivity or virulence.

Despite mounting evidence for the adaptive value of L452R, the exact functional or structural effect of the mutation and its impact on viral immunogenicity, pathogenesis, infectivity and/or transmissibility remains to be determined. Due to only recent attention to L452R variants, relatively few studies investigated the potential effects of this mutation. It was found that L452R reduces the Spike protein reactivity with a panel of the virus neutralizing antibodies and sera from convalescent patients [8, 14]. Moreover, it was found that out of 52 naturally occurring mutations in the receptor-binding motif residues of RBD that form the interface of direct interaction with ACE2, L452R results in the largest increase in free energy of the RBD-ACE2 binding complex, predicting stronger virus-cell attachment and, thus, increased infectivity [13]. We hope that the current and the original report on L452R variants of nCoV will facilitate in-depth structure-functional studies of the leucine residue in position 452 position and the change of hydrophobic leucine to a polar, highly hydrophilic arginine (or glutamine).

We cannot exclude the possibility of sample collection bias in our study as it was not originally designed as an in-depth surveillance study in specific geographical regions. In addition, our analysis is limited to a relatively small set of samples in hand and to publicly available genomes. However, we believe that the identification of CAL.20A and CAL.20C, with both common and unique features relative to the other circulating nCoV variant, will be useful in the optimization of real-time monitoring and the more complete understanding of the biological properties of this pandemic virus with a recently expanding number of genetic variations that are cause for significant public concern.

## Acknowledgement

We thank Prof. Steven Moseley for critical proofreading of the manuscript and scientific advice; clinical laboratory staff at Kaiser Permanente Washington and IEH Laboratories and Consulting Group for providing their support and technical expertise.

## METHODS

### Sample collection

Random de-identified nasopharyngeal samples were provided either as original swabs by Kaiser Permanente Washington (KPWA, April - May 2020, greater Seattle area) or as purified RNA by IEH Laboratories and Consulting group (September 2020 – February 2021, multiple states). Samples that tested positive for the presence of SARS-CoV-2 RNA (according to the respective laboratory practices) were subjected to further analysis. IEH samples of interest (see below) were assigned a unique identifier based on their collection date and source; information regarding the state of origin was provided for these samples for epidemiological analysis. Protected health information was completely removed from all clinical results before they were given to the researchers. All subjects cannot be identified directly or indirectly. The Western Institutional Review Board (Puyallup, WA) provided institutional biosafety committee services to Institute for Environmental Health by approving consent forms and human research safety protocols.

### RNA isolation and SARS-Cov-2 testing

RNA from KPWA samples was isolated using both AllPrep DNA/RNA Kit (Qiagen) and MagMax Viral Pathogen RNA isolation Kit (Thermo Fisher Scientific) according to manufacturer’s procedure. RNA isolation from IEH samples was performed according to laboratory guidelines using Thermofisher Kingfisher-96 instrument and proprietary IEH Nucleic Acid Extraction Reagent Kit. RNA was stored at −20oC until use. Testing of RNA for the presence of SARS-Cov-2 RNA at KPWA and IEH was performed according to laboratory guidelines using CDC 2019 Novel Coronavirus (2019-nCoV) Real-Time Reverse Transcriptase (RT)–PCR Diagnostic panel and proprietary IEH SARS-CoV-2 RT-PCR Test kit (based on the CDC RT-PCR kit), respectively.

### Amplification and sequencing of 414-583 and 1-250 regions

To amplify the 414-583 RBD region of the Spike gene from SARS-Coc-2 RNA samples, we used both one-step RT-PCR and two-step (cDNA synthesis followed by separate PCR amplification) reaction designs with commercial One-Step Ahead RT-PCR Kit (Qiagen) or IEH in-house RT, PCR and RT-PCR kits according to manufacturer’s guidelines. Both kits and conditions yield non-distinguishable results. Primers to amplify the region were as following: PF, 5’- GTGACATAGTGTAGGCAATGATG-3’, PR, 5’- TGGTGTAATGTCAAGAATCTCAAG-3’. First round of PCR (either RT-PCR on RNA or PCR on cDNA) consisted of 40 cycles of 10 sec 95oC, 15 sec 57oC, 40 sec 72oC. The product was then diluted 1:50 times with sterile water, and a second round of PCR was performed for 15 cycles at the same conditions using T7-tailed nested primers to obtain a single pure product (PF-T7Pro-nested, 5’- TAATACGACTCACTATAGGGCAAACTGGAAAGATTGC-3’, PR-T7Term-nested, 5’- GCTAGTTATTGCTCAGCGGCTCAAGTGTCTGTG-3’). Similarly, 1-250 Spike protein region was amplified firstly using primers PF2, 5’- CAGAGTTGTTATTTCTAGTGATGTTC-3’ and PR2, 5’- TGAAGAAGAATCACCAGGAGTC-3’, followed by the nested PCR on 1:50 diluted samples with tailed primers PF-T7Pro, 5’- TAATACGACTCACTATAGGGCAGAGTTGTTATTTCTAGTGATGTTC-3’, and PR-T7Term-nested2, 5’-GCTAGTTATTGCTCAGCGGGAGTCAAATAACTTC-3’. Sanger sequencing of the amplified region was performed from both ends by Eton Bioscience, Inc. Sequences were analyzed using BioEdit 7.2 and MEGA 7 Software.

### Whole genome sequencing

WGS was performed by IEH on MiSeq Illumina instrument; each sample was subjected to two individual rounds of sequencing. Sequences were assembled de novo using proprietary IEH pipeline or using the PATRICK sequence assembly service (https://patricbrc.org/app/Assembly2).

### Phylogenetic analysis of SARS-CoV-2 genomes

The latest global analysis of SARS-CoV-2 genomes from the Nextstrain database (https://nextstrain.org/ncov/global) was used to determine viral strains with the presence of L452 substitutions in the Spike protein (as of 12/17/2021). All genome sequences from PANGO lineages A.2.4, A.21, B.1.1.10, B.1.1.74. B.1.1.130, B.1.232, B.1.429, C.16 were downloaded from GISAID database (https://www.epicov.org/epi3, as of 12/17/2021). Sequences were checked for the presence of Spike-L452 substitution(s). All L452R-containing sequences submitted in January-February 2021 for lineages B.1.232 (CAL.20A) and 50 randomly chosen B.1.429 (CAL.20C) as well as the closely related L452 ancestors from both lineages were aligned and used to build total and synonymous-only phylogenetic trees using MEGA 7 software. Deletions of stretches of nucleotides which resulted in in-frame codon deletions were treated as single event. Unresolved nucleotides were assigned nucleotide value based on the closest relative. The same L452R sequences were used to run pairwise comparisons for synonymous substitutions for each CAL.20A and CAL20.C lineage separately and to calculate the average pairwise differences in silent mutations.

### Statistical analysis

Distribution of synonymous (S) and non-synonymous (NS) mutations within the 414-583 fragment was compared for RBD ridge epitope residues (aa 417, 443-452, 455-456, 470-490, 497—501, 570-572, total N = 49) vs other regions (N = 169) in McNemar test. There were 6 vs 1 NS and 7 and 8 S mutations, respectively, resulting in McNemar Chi2 4.50 (P =.034).

